# Polymer-based culture system enables expansion of human haematopoietic stem and progenitor cells while preserving ageing-associated transcriptional programs

**DOI:** 10.1101/2025.09.10.675303

**Authors:** Vasiliki Symeonidou, Laura Bond, Evangelia Giannelou, Yavor Bozhilov, Juan Rubio-Lara, Adam C. Wilkinson, George S. Vassiliou

## Abstract

The global rise in ageing populations is driving an increase in age-associated diseases, including haematological malignancies such as myelodysplastic syndromes and acute myeloid leukaemia. A major challenge in addressing this burden is the lack of experimental systems that enable mechanistic studies and therapeutic screening in ageing haematopoiesis. Here, we evaluate a polymer-based *ex vivo* culture platform for studying human haematopoietic stem and progenitor cells (HSPCs) in the context of ageing. Using CD34^+^ cells from cord blood, bone marrow and peripheral blood from different age groups, we show that this culture system enables robust expansion of HSPCs while preserving age-associated transcriptional programs. Single-cell RNA sequencing revealed enrichment of key progenitor populations with consistent transcriptional identities across donors. Importantly, expanded HSPCs from older individuals retained ageing-associated signatures, such as upregulation of inflammatory pathways. These findings demonstrate that our polymer-based expansion system enables robust modelling of ageing haematopoiesis and provide a single cell transcriptional landscape of this ageing model. We envision that this system could provide a versatile platform for mechanistic studies and pharmacological testing, addressing a critical gap in experimental frameworks for age-related haematological disease research.

## Introduction

The global population is ageing at an unprecedented rate, a demographic shift that is projected to intensify over the coming decades ^1^. Ageing is one of the most significant risk factors for disease, including cancer. Haematological malignancies exemplify this association as the incidence of conditions such as myelodysplastic syndromes and acute myeloid leukaemia rises markedly with advancing age ^2^. Consequently, the burden of haematological cancers is expected to grow substantially, presenting major clinical and healthcare challenges.

Ageing profoundly alters the haematopoietic system, leading to functional decline and increased susceptibility to disease. Advances in single-cell technologies have only recently enabled detailed characterisation of how ageing impacts haematopoiesis at the level of haematopoietic stem and progenitor cells (HSPCs) ^3–5^. Despite these extensive studies, a robust experimental framework for systematically evaluating genetic and pharmacological interventions in aged HSPCs is lacking. Such a framework requires an *ex vivo* system capable of expanding HSPCs without erasing their age-associated characteristics. Here, we investigated an *ex vivo* polymer-based human culture system recently developed for cord blood HSPCs in the context of HSPC ageing^6^. Remarkably, this system not only supports the expansion of adult HSPCs but also preserves their age-associated features, providing a platform for mechanistic studies and therapeutic screening in ageing haematopoiesis.

## Results

### Polymer-based culture system enables the expansion of adult bone marrow HSPCs

To initially assess whether the polymer-based culture system supports the expansion of adult HSPCs, we obtained CD34^+^ cells from donors across different age groups. Specifically, we collected CD34^+^ cells from two cord blood (CB) samples, two bone marrow (BM) samples from young adults (ages 32 and 34), and two BM samples from older adults (ages 60 and 72) (Figure 1A). The cells were cultured in expansion medium for two weeks according to the protocol described by Bozhilov *et al*., after which single-cell RNA sequencing was performed^6^.

**Figure 1.**
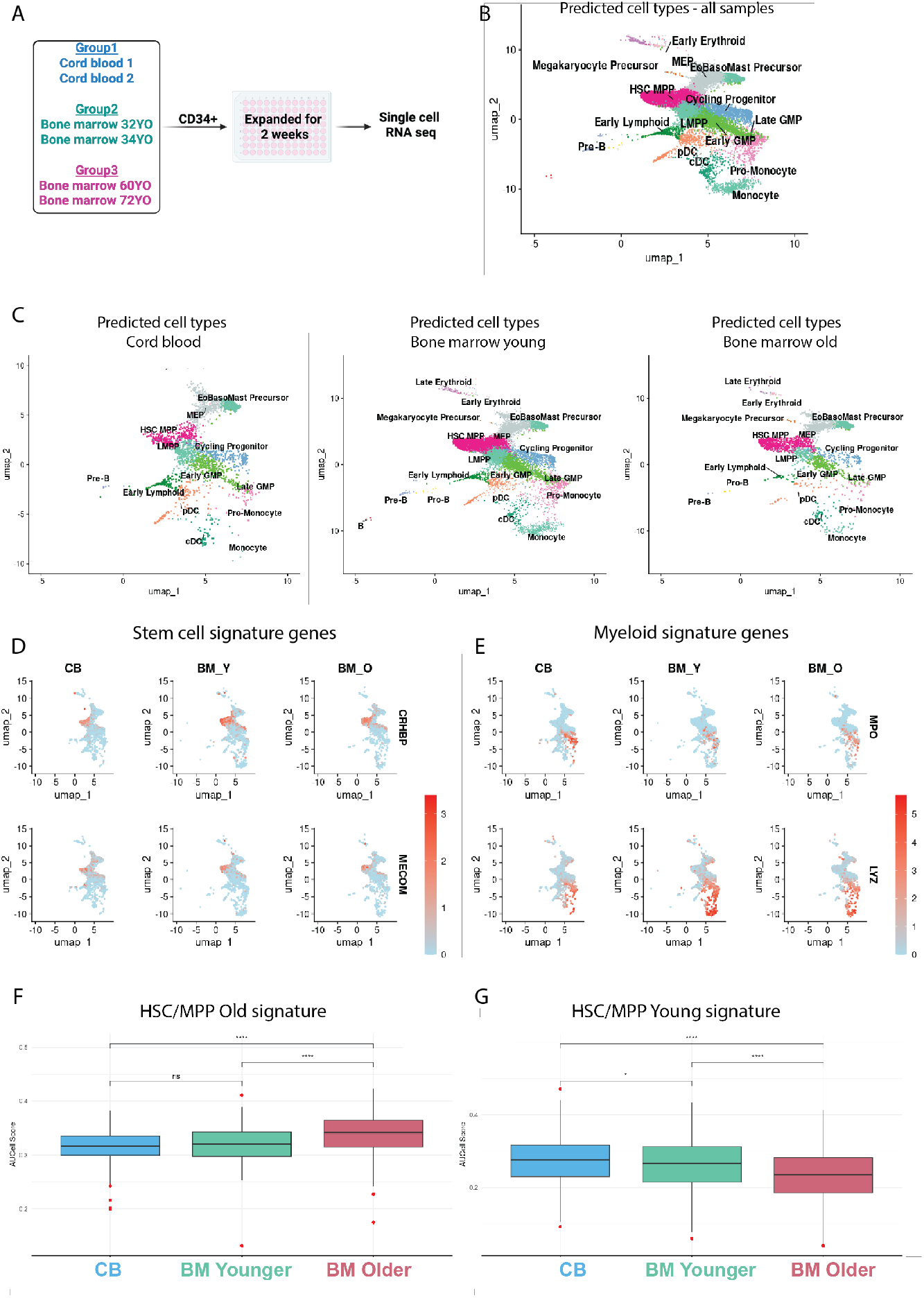
Polymer-based culture system enables the expansion of adult bone marrow HSPCs. A) Experimental outline. Created with BioRender.com B) UMAP of predicted cell types from scRNA-seq - all samples combined. C) UMAP of predicted cell types by HSPC sample type/source (cord blood, bone marrow young, bone marrow old). D) Expression of stem cell signature genes *CRHBP* and *MECOM* in different HSPC sample types. E) Expression of myeloid signature genes *MPO* and *LYZ*. F) AUCell score for “old” signature. G) AUCell score for “young” signature. Significant differences were determined using the Wilcoxon rank-sum test with Benjamini–Hochberg correction. **** *p*.*adj* < 0.0001

To explore the cell-types present in these HSPC cultures, we mapped our single-cell RNA sequencing data to a single-cell reference atlas of human hematopoietic differentiation (BoneMarrowMap)^7^. Predicted cell types were consistent across all samples, and clustering patterns were uniform across the different age groups (Figure 1B). In all three age groups, we observed enrichment of hematopoietic stem and progenitor populations, including hematopoietic stem cells and multipotent progenitors (HSC/MPP cluster), lymphoid-primed multipotent progenitors (LMPPs), and cycling progenitors. Additionally, we detected expansion of megakaryocyte–erythroid progenitors (MEPs) and eosinophil– basophil–mast precursors (EoBasoMast) (Figure 1C and Supplementary Figure 1A).

To validate the transcriptional profiles of each cluster, we examined specific gene signatures. *CRHBP* ^*8*^ and *MECOM* were used as stem cell markers (Figure 1D), *MPO* and *LYZ* as myeloid markers (Figure 1E), and *ITGA2B* and *GATA2* as erythroid/megakaryocytic markers (Supplementary Figure 1B). The observed expression patterns matched the predicted cell types, providing confidence in our cluster assignments.

We next investigated whether the expansion culture preserves ageing-associated transcriptional programs. To assess this, we focused on the HSC/MPP cluster and compared our data with age-associated HSC transcriptional signatures from Jakobsen *et al*.^*9*^. AUCell scores revealed increased activity of ageing-associated transcriptional programs in the HSC/MPP cluster of older individuals compared to young donors (Figure 1F). Conversely, HSC/MPPs from younger donors showed stronger activity of young-associated transcriptional programs (Figure 1G).

To further investigate transcriptional differences between old and young BM–expanded HSC/MPPs, we performed differential expression analysis followed by Gene Set Enrichment Analysis (GSEA)^10^. The top upregulated pathways in old BM HSPCs were Interferon Gamma Response and Interferon Alpha Response (Supplementary Figure 1C). Upregulation of these pathways is a hallmark of ageing^5^, suggesting that the expansion system preserves intrinsic states such as inflammation^11^. These results demonstrate that the polymer-based culture system supports the expansion of HSPCs from both young and aged BM while preserving age-associated transcriptional programs.

### Polymer-based culture system enables the expansion of adult peripheral blood HSPCs

As bone marrow samples are difficult to obtain, we sought alternative and more readily available sources of CD34^+^ cells. One such source is leukapheresis cones, which are by-products of routine leukodepletion procedures. These cones contain large numbers of peripheral blood mononuclear cells, including CD34^+^ HSPCs, making them a widely available and cost-effective resource for research.

As before, we obtained CD34^+^ cells from donors across different age groups and cultured them in our expansion system (Figure 2A). Mirroring our observations from BM samples, we detected enrichment in the HSPC clusters (Figure 2B–C and Supplementary Figure 1A). To confirm reproducibility, we repeated the experiment with an independent donor and again observed *ex vivo* expansion of HSPCs (Supplementary Figure 2A–B and 1A). To test whether ageing-associated transcriptional programs are preserved in peripheral blood–derived expanded cells, we performed AUCell analysis as before. We found higher activity of ageing-associated programs in the HSC/MPP cluster of older compared to younger donors (Figure 2D). Conversely, HSC/MPPs from younger donors showed stronger activity of young-associated programs (Figure 2E). We also investigated the expansion rate between HSPCs from different age groups, and we observed a higher expansion rate in CB-derived cells compared to cells from adult donors (Supplementary Figure 2C).

**Figure 2.**
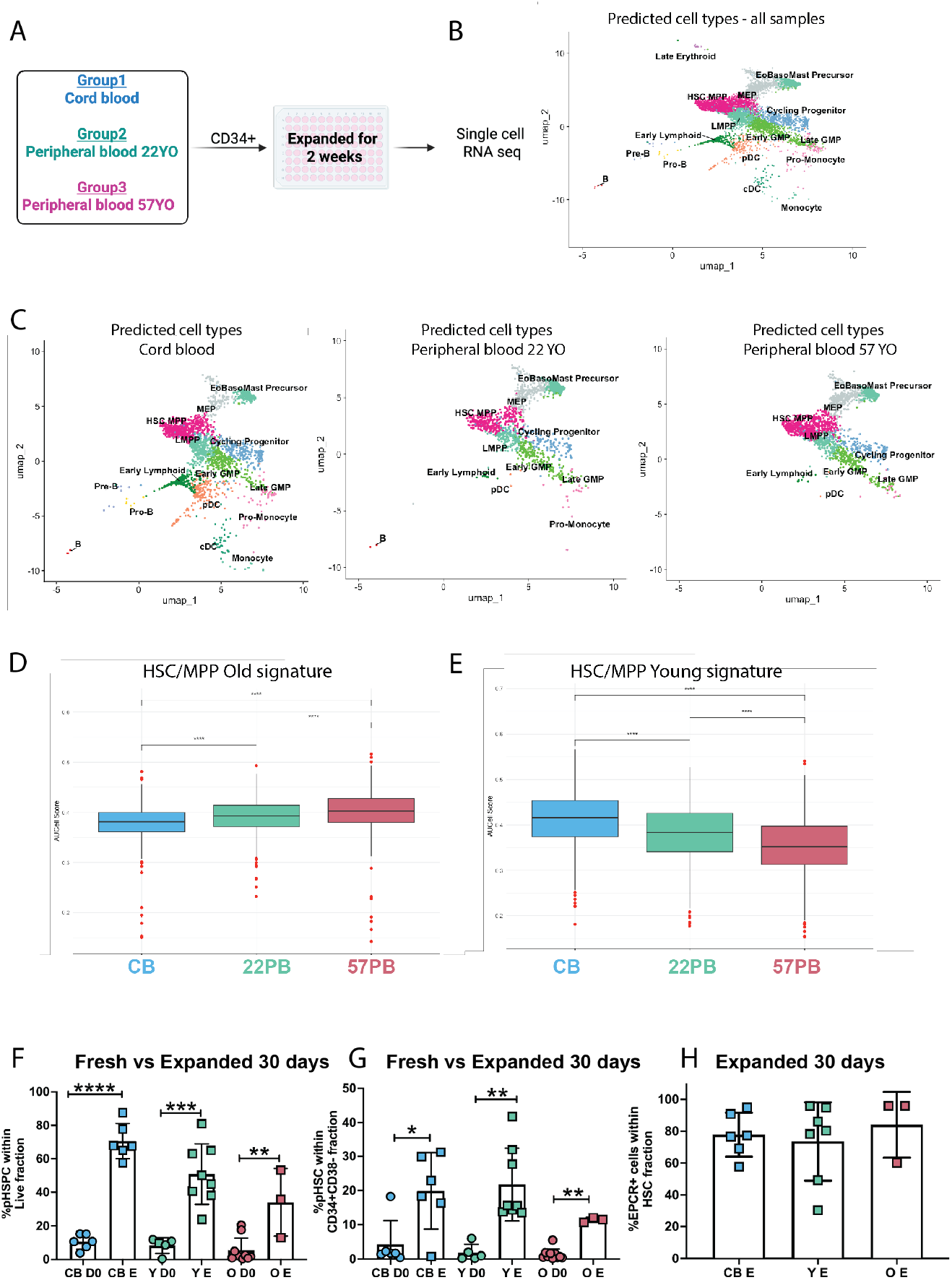
Polymer-based culture system enables the expansion of adult peripheral blood HSPCs. A) Experimental outline. Created with BioRender.com B) UMAP of predicted cell types from scRNA – all samples combined. C) UMAP of predicted cell types by HSPC sample type/source (cord blood, peripheral blood young, peripheral blood old). D) AUCell score for “old” signature. E) AUCell score for “young” signature. Significant differences were determined using the Wilcoxon rank-sum test with Benjamini– Hochberg correction. **** *p*.*adj* < 0.0001. F) % phenotypic haematopoietic stem and progenitor cells (pHSPCs) within the live fraction. Data are shown as mean +/-SD. Mann-Whitney test was performed. G) % phenotypic haematopoietic stem cells (pHSCs) within the CD34^+^CD38-fraction. Data are shown as mean +/-SD. Mann-Whitney test was performed. H) % of EPCR+ cells within the HSC fraction from cells expanded for 30 days. \**p* < 0.05, \*\**p* < 0.01, \*\*\**p* <0.001, \*\*\*\**p* < 0.0001.

To test whether the culture supports long-term HSPC expansion, we cultured CB and peripheral blood-derived CD34^+^ cells for 30 days. At the end of the culture, we performed immunophenotyping to compare expanded cells with freshly isolated CD34^+^ cells. We observed an increase in the percentage of both the immunophenotypic HSPC and HSC fractions relative to fresh CD34^+^ cells (Figure 2F-G). Since CD38 is not expressed in cultured cells, we further examined EPCR (CD201) expression as a marker of HSC identity^12^ and found that the majority of HSCs in our culture were EPCR+(Figure 2H). These results demonstrate that leukapheresis cones provide a reliable and reproducible source of HSPCs that can be expanded *ex vivo* while preserving intrinsic age-associated transcriptional signatures.

## Discussion

Understanding how ageing impacts human haematopoiesis is crucial for deciphering the increased susceptibility to haematological malignancies of older individuals. Our study demonstrates that the polymer-based *ex vivo* culture system developed by Bozhilov *et al*. supports the robust expansion of HSPCs from multiple sources, including cord blood, bone marrow, and peripheral blood^6^. Notably, the system preserves age-associated transcriptional programs, allowing the intrinsic differences between young and old HSC/MPPs to remain detectable after expansion.

Peripheral blood-derived CD34^+^ cells from leukapheresis cones proved to be an accessible and reproducible source of HSPCs. Immunophenotyping confirmed that the majority of expanded HSCs were EPCR+, consistent with a *bona fide* stem cell identity, even after long-term culture. The ability to maintain EPCR expression alongside age-associated transcriptional signatures is particularly important, as it ensures that expanded cells retain both functional and molecular hallmarks of their *in vivo* counterparts.

Our findings demonstrate that the polymer-based culture system developed by Bozhilov *et al*., is a suitable platform for use in systematic investigation of age-associated functional decline and its consequences in human HSPCs. By combining accessible sample sources with an expansion system that preserves intrinsic donor characteristics, this platform enables mechanistic studies and pharmacological testing in a physiologically relevant context. Future work could leverage this system to identify interventions that mitigate age-related haematopoietic dysfunction or target ageing-specific vulnerabilities in haematological malignancies and pre-malignancies.

**Supplementary Figure 1.**
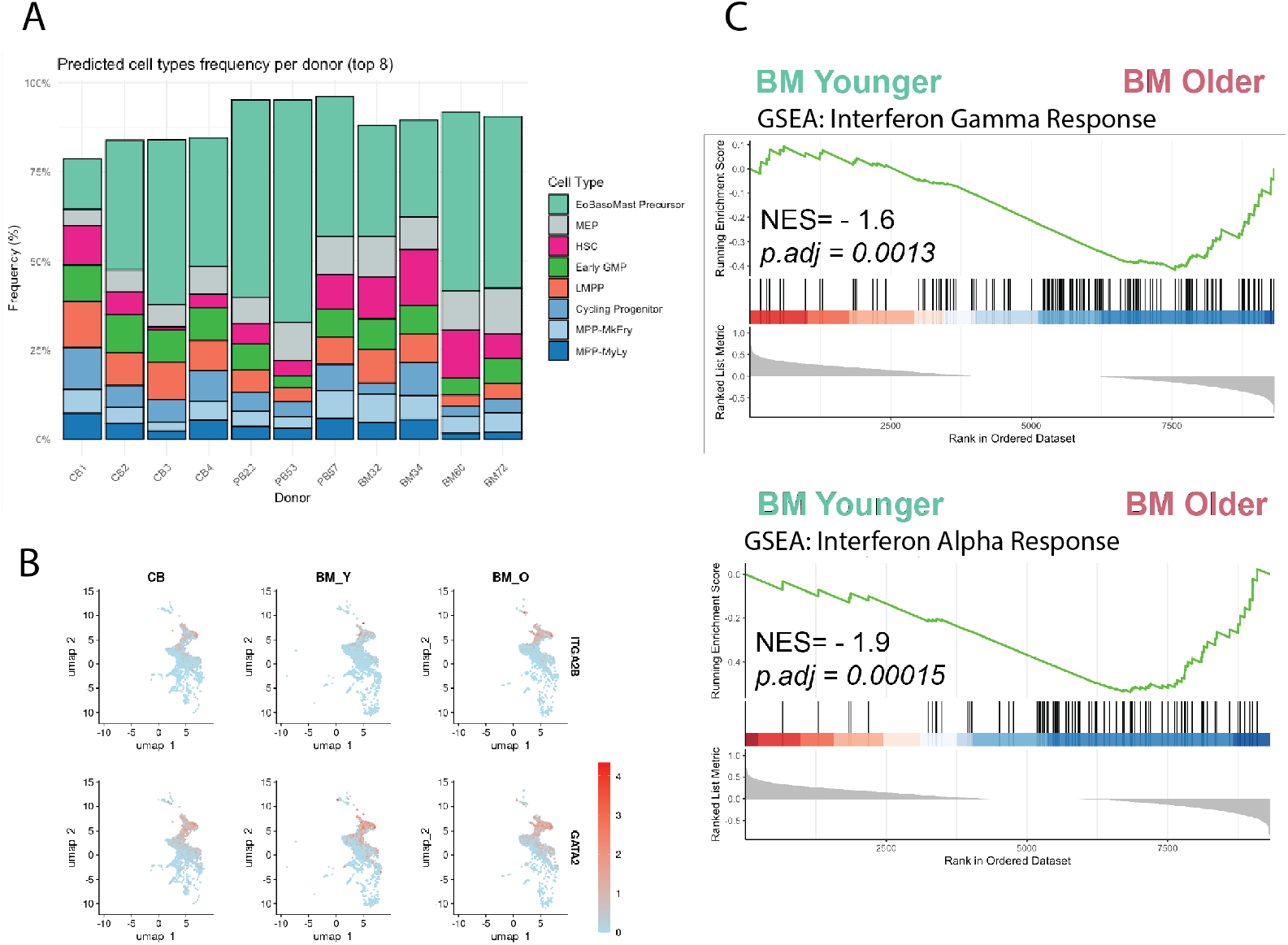
A) Frequency of top 8 predicted cell types across all samples. B) Expression of erythroid/megakaryocytic genes. C) Gene Set Enrichment Analysis identified upregulation of Interferon Gamma Response and Interferon Alpha Response gene sets in expanded cells from bone marrow from older donors.

**Supplementary Figure 2.**
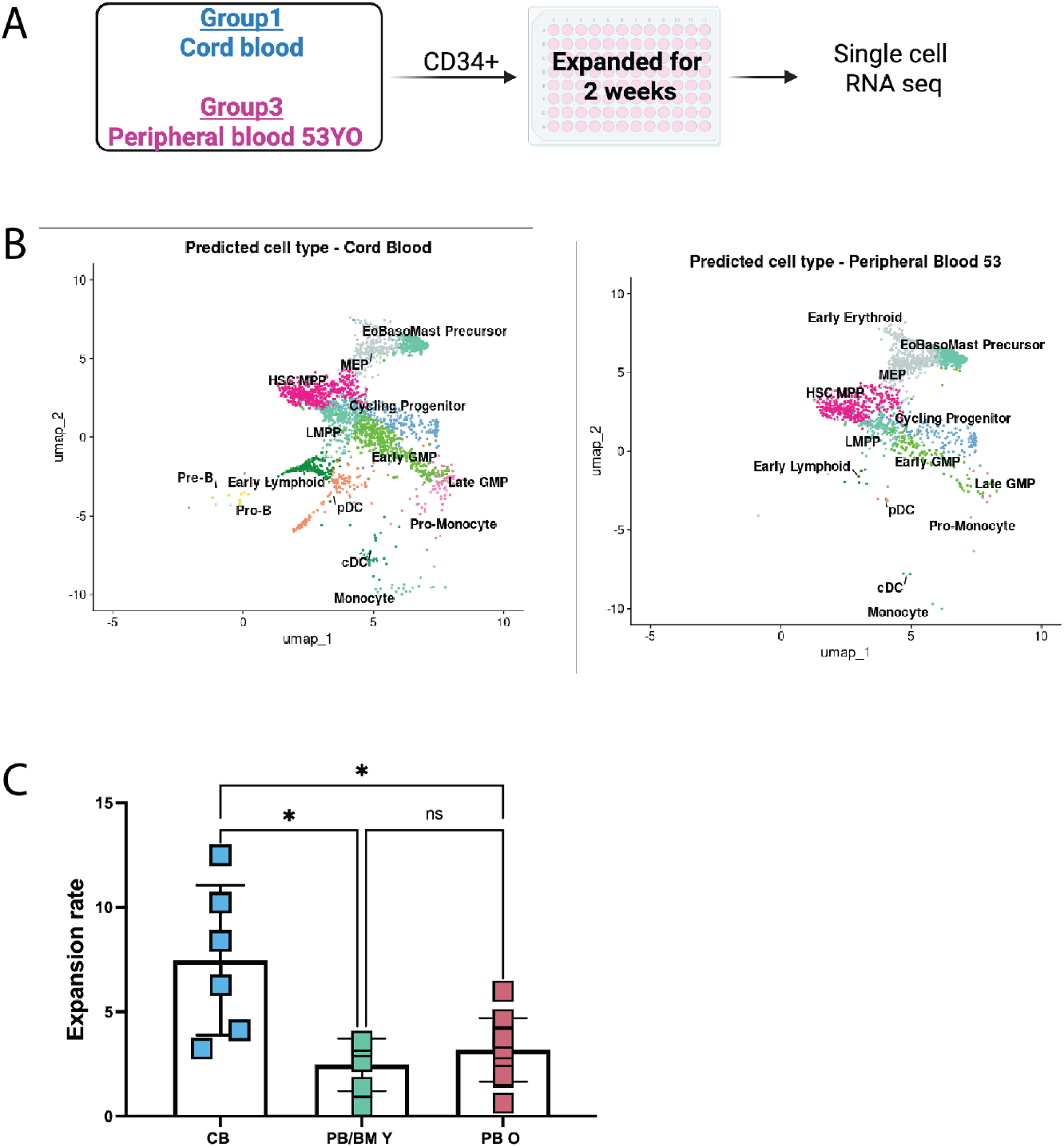
A) Experimental plan. Created with BioRender.com B) UMAP of predicted cell types by HSPC sample type/source (cord blood and peripheral blood from 53-year-old. C) Expansion rate after 14 days. Data are shown as mean +/-SD. 2-way Anova test was performed. \**p* < 0.05,

## Materials and Methods

### Human sample sources and ethics

Leukocyte cone samples were purchased from NHS Blood and Transplant (NHSBT). These were collected fresh from platelet apheresis donors in the Cambridge Blood Donor Centre. Cord blood samples were purchased from NHSBT. Fresh samples were collected from NHSBT sites and sent via courier, within 24 hours. Bone marrow hip samples were collected from individuals undergoing elective total hip replacement (THR) surgery at the Addenbrookes Cambridge University Hospital. Informed consent was received from all participants prior to surgery. All samples were obtained through the Cambridge Blood and Stem Cell Biobank (CBSB). This study was approved by the East of England - Cambridge East Research Ethics Committee (REC Ref: 24/EE/0116).

### Sample collection and processing

Leukocyte cones: Leukocyte cones are filters from the Leukocyte Reduction System (LRS) used during platelet donation to capture white blood cells. All samples were processed within 24 hours of collection. After initially drip-draining the cells out from the leukocyte cone, samples were diluted 1:1 with PBS/0.5% BSA (Phosphate Buffered Saline with 0.5% Bovine Serum Albumin). Mononuclear cells were isolated using Lymphoprep density gradient centrifugation (StemCell Technologies).

Cord blood samples: Fresh cord blood samples were collected into a collection bag. All samples were processed within 48 hours of collection. Samples were poured out of the collection bag port and then diluted 1:1 with PBS/0.5% BSA. Mononuclear cells were isolated using Lymphoprep density gradient centrifugation (StemCell Technologies).

Bone marrow hip samples: From each THR patient, the femoral head was retained. Additionally, a bone marrow aspirate sample was obtained from the femoral canal and collected into an 8.2ml 9NC S-Monovette tube. All samples were processed within 24 hours of collection. The bone marrow aspirate was filtered through a 70 µm cell strainer to remove any remaining clots and bone fragments. The sample was then diluted 1:1 with PBS/0.5% BSA. Mononuclear cells were isolated using Lymphoprep density gradient centrifugation and a SepMate tube (StemCell Technologies). Bone was manually extracted from the exposed base of the femoral head (including the femoral neck where present) using a flat, straight end metal spatula. The extracted bone was then manually fragmented using the same spatula. The fragments were thoroughly washed with PBS/0.5% BSA and filtered through a 70 µm cell strainer. The sample was then centrifuged (1800rpm for 5 minutes) and the cell pellet transferred to a new tube to reduce the presence of oil/adipocytes in the sample.

CD34^+^ cell selection: All samples were then enriched for CD34^+^ cells using a CD34 MicroBead Kit UltraPure and an AutoMACS machine (Miltenyi Biotec), according to the manufacturer’s protocols. The CD34^+^ enriched fraction was then frozen in 90% KnockOut Serum Replacement (KOSR, Gibco) with 10% dimethyl sulfoxide (DMSO). All samples were stored in liquid nitrogen until use in downstream experimentation.

### Single cell RNA sequencing library preparation

Single-cell suspensions from each biological sample were stained with unique Cell Multiplexing Oligos (CMOs) using the Chromium CellPlex kit (10x Genomics). Briefly, up to 1× 10^6 cells per sample were pelleted at 300–400 × g for 5 min at 4 °C, resuspended in the manufacturer’s CMO labeling buffer, and incubated with the assigned CMO at the recommended working concentration for 5 min at room temperature with gentle mixing. Labelling was quenched and 3 washes in PBS + 1% BSA were performed to remove unbound CMO. Cell were then sorted in BD Aria Fusion.

Gel beads-in-emulsion (GEMs) were generated on a Chromium Controller (10x Genomics) using Chromium Next GEM Single Cell 3’ reagents (per manufacturer protocols). The pooled cells, Master Mix, Partitioning Oil, and gel beads were loaded as directed. Reverse transcription (RT) was performed in-GEM to barcode polyadenylated transcripts with 10x barcodes and UMIs. Following RT, emulsions were broken and cDNA was recovered and cleaned up with silane and SPRI bead workflows supplied in the kit. Full-length barcoded cDNA was PCR-amplified (cycles adjusted per the manufacturer’s guidelines and the cDNA yield of each run), purified with SPRI beads, and quality-checked by Agilent Bioanalyzer and Qubit. Gene Expression libraries were constructed from amplified cDNA using the 10x Genomics 3’ library prep reagents. CellPlex CMO libraries were prepared in parallel from the same cDNA according to the CellPlex protocol. Final libraries were quantified by qPCR and profiled for size distribution.

Gene Expression and CMO libraries were pooled at recommended ratios and sequenced on an Illumina NovaSeq using paired-end chemistry with standard 10x read structure for 3’ Gene Expression and CellPlex. Read 1 captured the cell barcode and UMI; Read 2 captured the transcript insert; index reads captured sample and CMO indices. Read lengths followed 10x Genomics recommendations (R2 ≥ 90–100 bp for robust mapping). Libraries were sequenced to a depth of ~[20,000–50,000] read pairs per cell for Gene Expression and ~[2,000–5,000] reads per cell for CMO libraries, adjusted to the complexity of each experiment.

### Flow cytometry

Flow cytometry was performed of BD FACS using BD Aria Fusion. Antibodies used CD34 (clone 581) CD38 (clone HIT2), CD45RA(HI100), CD90 (5E10), CD49f (GoH3), EPCR (RCR-401), CD123 (clone 6H6), Lineage markers include: CD2 (RPA-2.10), CD3 (SK7), CD7 (CD7-6B7), CD11b (ICRF44), CD14 (61D3), CD19 (HIB19), CD20 (2H7), CD56 (NCAM16.2), CD235a (HIR2), CD41a (HIP8) for viability marker eBioscience Fixable Viability Dye eFluor 780. Briefly, cells were washed twice with PBS, followed by incubation with a viability dye on ice for 20 minutes. After incubation, the cells were washed with PBS containing 2% FBS. Finally, the cells were stained in BD Brilliant Stain Buffer for 30 minutes on ice. Flow cytometry data were analysed using FlowJo 10.

### Bioinformatics Analysis

Samples were processed using the 10x Genomics pipeline. Briefly, sequenced scRNA-seq libraries were demultiplexed and processed using the CellRanger Single-Cell Software Suite (v.9.0.0 10x Genomics) using GRCh38 (hg38) human reference genome. Unique molecular identifiers (UMIs) were quantified to generate a cell-by-gene count matrix for each sample. These matrices were then imported into the R environment (v.4.4.1) and analysed using Seurat (v.5.3.0). RNA counts were log-normalized and filtered (500–7,500 features; <5% mitochondrial). CMO counts were CLR-normalized, demultiplexed with HTODemux, and singlets retained. QC and visualization were performed using standard Seurat functions. Datasets from CMO experiments were loaded as Seurat objects and merged. Counts were normalized using SCTransform. Pre-integration batch effects were visualized, and batch correction was performed using Harmony^13^, with UMAP recomputed on Harmony embeddings. The integrated dataset was projected onto the BoneMarrowMap^7^ using map_Query for hematopoietic cell type annotation. Cells failing mapping QC were excluded. Hematopoietic pseudotime and cell types were predicted using a K-nearest neighbors (KNN) approach, and UMAPs were generated for visualization. Gene signatures for aging were defined based on Jakobset *et al*.^*8*^. Expression matrices were extracted from the equalised Seurat object, cell-level rankings computed with AUCell_buildRankings(), and gene set activity quantified using AUCell_calcAUC() (aucMaxRank = 0.05 × nGenes) using AUCell^14^. AUC scores were merged with metadata for visualization. Differential expression was performed using Seurat’s FindMarkers(), and significant genes were used for downstream gene set enrichment analysis (GSEA)^10^.

### Statistical analysis

Data analysis and statistical tests were performed using R version v.4.4.1and GraphPad Prism v10.5.0.

## Author contributions

V.S. conceived and designed the study, performed the bioinformatic analysis and the majority of the experiments and wrote the manuscript. L.B. performed additional experimental work. E.G. assisted in obtaining donor material. Y.B. and J.L.R. and A.C.W. provided technical advice and expertise. G.S.V. supervised the study. A.C.W. and G.S.V. contributed to writing the manuscript. All authors reviewed the final paper.

